# LRP5-deficiency in OsxCreERT2 mice models intervertebral disc degeneration by aging and compression

**DOI:** 10.1101/720656

**Authors:** Matthew J. Silva, Nilsson Holguin

## Abstract

Osterix is a critical transcription factor of mesenchymal stem cell fate, where its loss or loss of WNT signaling diverts differentiation to a chondrocytic lineage. Intervertebral disc (IVD) degeneration activates differentiation of prehypertrophic chondrocyte-like cells and inactivates WNT signaling, but its interacting role with osterix is unclear. First, compared to young-adult (5mo), mechanical compression of old (18mo) IVD induced greater IVD degeneration. Aging (5 vs 12mo) and/or compression reduced the transcription of osterix and notochordal marker T by 40-75%. Compression elevated transcription of hypertrophic chondrocyte marker MMP13 and pre-osterix transcription factor RUNX2, but less so in 12mo IVD. Next, using an Ai9/td reporter and immunohistochemistry, annulus fibrosus and nucleus pulposus cells of 5mo IVD expressed osterix, but aging and compression reduced its expression. Lastly, in vivo LRP5-deficiency in osterix-expressing cells degenerated the IVD, inactivated WNT signaling, reduced the biomechanical properties by 45-70%, and reduced transcription of osterix, notochordal markers and chondrocytic markers by 60-80%. Overall, these data indicate that age-related inactivation of WNT signaling in osterix-expressing cells may limit regeneration by depleting progenitors and attenuating the expansion of chondrocyte-like cells.

## INTRODUCTION

Intervertebral disc (IVD) degeneration is a multi-faceted disease physically characterized by dehydration, height loss and, in the later stages, calcification (Boos et al., 1997; Rutges et al., 2010) and annulus fibrosus (AF) rupture. Aging and IVD degeneration lead to degradation of the extracellular matrix (Antoniou et al., 1996), potentially from a shift in the population of resident cells in the IVD (Hunter et al., 2004), cell loss (Hunter et al., 2004; Urban and Roberts, 2003), altered cell metabolism, among other changes. Chondrocyte-like cells of the IVD resemble articular chondrocytes in their appearance and transcriptional expression and cohabitate with notochordal cells in the nucleus pulposus (NP) (Clouet et al., 2009; Minogue et al., 2010), but are smaller and phenotypically distinct from notochordal cells (Chen et al., 2006; Minogue et al., 2010). Aging and IVD degeneration induce the disappearance of notochordal cells (Richardson et al., 2017), which are replaced by chondrocyte-like cells (Boos et al., 2002; Yurube et al., 2014), ostensibly via differentiation of prehypertrophic chondrocyte-like cells (Rutges et al., 2010). RUNX2 (Cbfa1), Sp7 (Osterix) and Ctnnb1 (β-Catenin) progressively drive skeletal progenitors to become osteoblasts and later osteocytes, but loss of osterix or WNT signaling diverts cell fate towards chondrogenesis (Long, 2011). Previously, we showed that mediating WNT signaling impacts notochordal expression in the IVD (Holguin and Silva, 2018), but it is unknown if WNT signaling directly impacts transcription factor osterix.

Transcription factor β-Catenin is regulated by the canonical WNT pathway (Milat and Ng, 2009) and is putatively involved in the regeneration of IVDs. Patients with IVD degeneration have up-regulated levels of b-Catenin (Wang et al., 2012). In vivo, β-Catenin is promoted in canine intervertebral discs with age-related propensity for degeneration (Smolders et al., 2012). Aging inactivates WNT signaling and disrupting WNT/β-Catenin signaling during development deteriorates the entire IVD (Kondo et al., 2011; Mundy et al., 2011). Re-activating WNT signaling in IVDs of aged mice leads to greater aggrecan (Winkler et al., 2014), but greater β-Catenin in rodent IVD cells also triggers cellular senescence, apoptosis and biomarkers of matrix breakdown (Hiyama et al., 2010).

Here, we apply in vivo chronic loads to a range of adult IVD to demonstrate transcriptional downregulation of osterix in IVD degeneration by aging and mechanical compression. Further, we show that aging and compression reduce the protein expression of osterix in the annulus fibrosus and nucleus pulposus of the IVD. Lastly, targeted suppression of WNT signaling in the cells of the IVD that express osterix (using an OsxCreERT2 driver) induces IVD degeneration to a degree similar to aging and compression, as demonstrated by histology, qpcr and biomechanics. Together, these data suggest that limited WNT signaling in older IVD may potentiate IVD degeneration by attenuating the expansion of chondrocyte-like cells.

## METHODS

### Mice

Female TOPGAL (TCF/LEF Optimal Promoter/Galactosidase reporter) transgenic mice that were aged to 5 (n=7) or 12 months (n=6) from a previous experiment were used to determine mRNA expression changes with tail compression (Holguin and Silva, 2018). Female C57Bl/6J mice were purchased from the National Institute of Aging (NIA, Bethesda, MD, USA) (5 mo: n=8, 18 mo: n=5 and 22 mo: n=l). A separate set of female C57Bl/6J mice (n=3) were tail compressed to stain by IHC for osterix protein. To generate Osx-CreERT2/LRP5^fl/fl^/TOPGAL/tdT mice (LRP5 cKO, n=6), OsxCreERT2 mice were crossed with LRP5^fl/fl^ mice (receptor of WNT signaling), TOPGAL mice (reporter of WNT signaling) and Ai9(RCL-tdT) (fluorescent reporter of location of osterix) mice. Wildtype mice were LRP5^fl/fl^/TOPGAL/tdT (WT, n=6). To suppress WNT signaling, LRP5 cKO mice were provided tamoxifen in their chow (Envigo, Indianapolis, IN) for 5 days per week for 1 month; WT mice also received tamoxifen chow. To report osterix location, mice were injected with tamoxifen for two days and the intervertebral discs harvested on the third day. WT mice (No Cre) served as controls. All mice were housed 4-5 per cage under standard conditions with ad libitum access to water and either Tamoxifen chow as noted above, or regular chow (Purina 5053 & 5058, Purina, St. Louis, MO). The study was approved by the Washington University Animal Studies Committee.

### Tail Compression

Once the CC7 (Caudal Coccygeal 7) and CC9 vertebra were identified with preoperative radiographs, 23-G needles were implanted transcutaneously during anesthesia by isoflurane (2.5% vol) and postoperative radiographs confirmed proper placement. Compression rings were attached to the pins to apply mechanical load via tightening of four screws with compressive springs (Cat # 9001T24, McMaster, Elmhurst, IL). Following injection with Buprenorphine (1 mg/kg s.c.) for pain relief, 2.25 N of load was applied for 1 week to induce degeneration. As a positive control for severe IVD degeneration, 5 mo tail IVD were punctured (n=3).

### MicroCT

CC6-CC7 motion segments were imaged by micro-computed tomography (VivaCT 40, Scanco Brüttisellen, Switzerland) at a resolution of 21 μm (70 kV, 114 μA, and 100 ms integration time) to determine the IVD morphology and normalize the mechanical properties. Semi-automatic contouring was used to segment the IVD from bone with a lower/upper threshold of 205/1,000 (406 mg HA/cm^3^).

### Mechanical Testing

Controlled mechanical tests were performed on CC6-CC7 motion segments as previously described (Holguin et al., 2013). Prior to mechanical testing, the bone-disc-bone segments were excised, zygapophysial joints and superficial tissue were removed, and spinal units were hydrated in 1X PBS for 18 h at 4 °C. Superior and inferior vertebra were gripped by microvises and, once secured, the sample was immersed in 1X PBS. A materials testing system (Electropulse 1000, Instron, Norwood, MA, USA) applied twenty compression-tension cycles at a frequency of 0.5 Hz and the limit under displacement-control was determined by noting linear stiffnesses.

### Mechanical Data Analysis of IVDs

A trilinear fit model determined the compressive, tensile and neutral zone stiffness of the motion segment. Briefly, the compressive and tensile loading curves were isolated and a 6th order polynomial was fit to the 20th loading and unloading tension- compression cycle. The minimum derivative of the curve represented the neutral zone stiffness and the derivative measured at 80% of the maximum load magnitude in the compressive and tensile portion of the curve constituted the compressive and tensile stiffness, respectively. The material properties (moduli) were determined by multiplying stiffness by the height of the IVD and dividing by the area.

### Histology, Immunohistochemistry and Frozen Sectioning

Beta-galactosidase staining for WNT activity, Safranin-O, and IHC was performed as previously described (Holguin et al., 2016) on lumbar (L1-L3) and coccygeal (CC8-CC10) motion segments. Motion segments were freshly harvested, fixed in 4% paraformaldehyde for 1 h, incubated in Xgal (Invitrogen, Grand Island, NY) for 48 h, fixed overnight, decalcified with Immunocal for two days and embedded in paraffin using routine methods. These, coronal sections (10 μm) were used to visualize galactosidase cells and serial sections were counter stained with eosin or Safranin-O. All other intervertebral discs from other mice were not incubated with Xgal or fixed overnight. Sections (5 μm) for immunohistochemistry were deparaffinized and stained with osterix antibody (#22552, Abcam) and counter stained with Alcian blue. Cytomorphology will be used to distinguish notochordal and chondrocyte-like cells (Hunter et al., 2004; Smolders et al., 2012). For frozen section tissues were fixed in 4% paraformaldehyde, decalcified in 14% EDTA for 3 days, infiltrated with 30% sucrose overnight, embedded in OCT, sectioned coronally (10 μm), and stained with DAPI.

### Analysis of Histology and qPCR

Histological scoring (Tam et al., 2019) was accomplished using Safranin-O/Fast green images of IVDs and was on a 14-point scale. In brief, the nucleus pulposus, annulus fibrosus and boundary between the nucleus pulposus and annulus fibrosus of the IVD were scored and added together for a total IVD score. WNT activity (LacZ expression) quantification within the nucleus pulposus and annulus fibrosus were carried out as previously described (Holguin et al., 2014). For instrumented animals, tail IVDs between CC7-9 and CC10-12 and, for genetic mouse models, tail IVDs CC10-12 and lumbar discs L3-5 were separated from all other tissues and snap frozen in liquid nitrogen. Then samples were pulverized in a Mikro Dismembrator (B. Braun Biotech International Mikro-Dismembrator S, Germany) and suspended in TRIZOL (Ambion) until further processing. Total RNA extraction was performed using a standard kit (RNeasy mini kit, Qiagen). RNA concentration was quantified (ND-1000, Nanodrop). First strand cDNA was synthesized (iScript, Biorad) from 500 ng of total RNA for Taqman (Life Technologies) probes. The relative gene expression in loaded and control IVDs was determined by normalizing to housekeeping gene *IPO8* (Mm01255158_m1) and then normalized to the control IVD (2^−ΔΔCT^) or, in KOs, to the average WT value.

### Statistics

An ANOVA with Bonferroni post hoc test was used to compare histological scoring of 5 mo and 18 mo IVD subjected to mechanical injury. A two way ANOVA was used to compare qPCR and osterix protein expression (dark vs light vs no stain) with loading (Control vs Loaded). Unpaired Student’s t-tests compared intervertebral discs of WT to genetic KO animals. Linear regression correlated the relative expression (cKO/WT) of WNT signaling genes to b-Catenin, tensile stiffness, RUNX2 and aggrecan. Statistical computations were completed using SPSS (IBM SPSS Statistics 25) and significance was set at p<0.05.

## RESULTS

### Aging and mechanical compression induce IVD degeneration

In order to determine the regulation of osterix by IVD degeneration, a gradation of IVD degeneration was created by subjecting mice of different adult ages to mechanical loading (compression). Aging increased the IVD degeneration of lumbar (**Fig. 1A**) and tail IVD (**Fig. 1B, C**). Notable changes in 15-18 mo IVD included loss of proteoglycan (red staining), cell death, disruption of the NP cell band and large rounded chondrocytes in the inner AF. Further, aging in the lumbar IVD (22 mo), included calcification of the NP, cell cloning (cell clusters) and loss of the demarcation between the NP and AF. Mechanical compression of the young-adult IVD induced scalloping and reversal of the inner AF. Further, mechanical loading of aged IVD induced severe proteoglycan loss, calcification of the IVD and loss of the demarcation between the NP and AF. Puncture of the IVD induced most of the above-mentioned features of IVD degeneration and fissures in the AF. Taken together, aging and mechanical injury each induced IVD degeneration and together had an additive effect, but the underlying mechanism was unclear.

**Figure 1.**
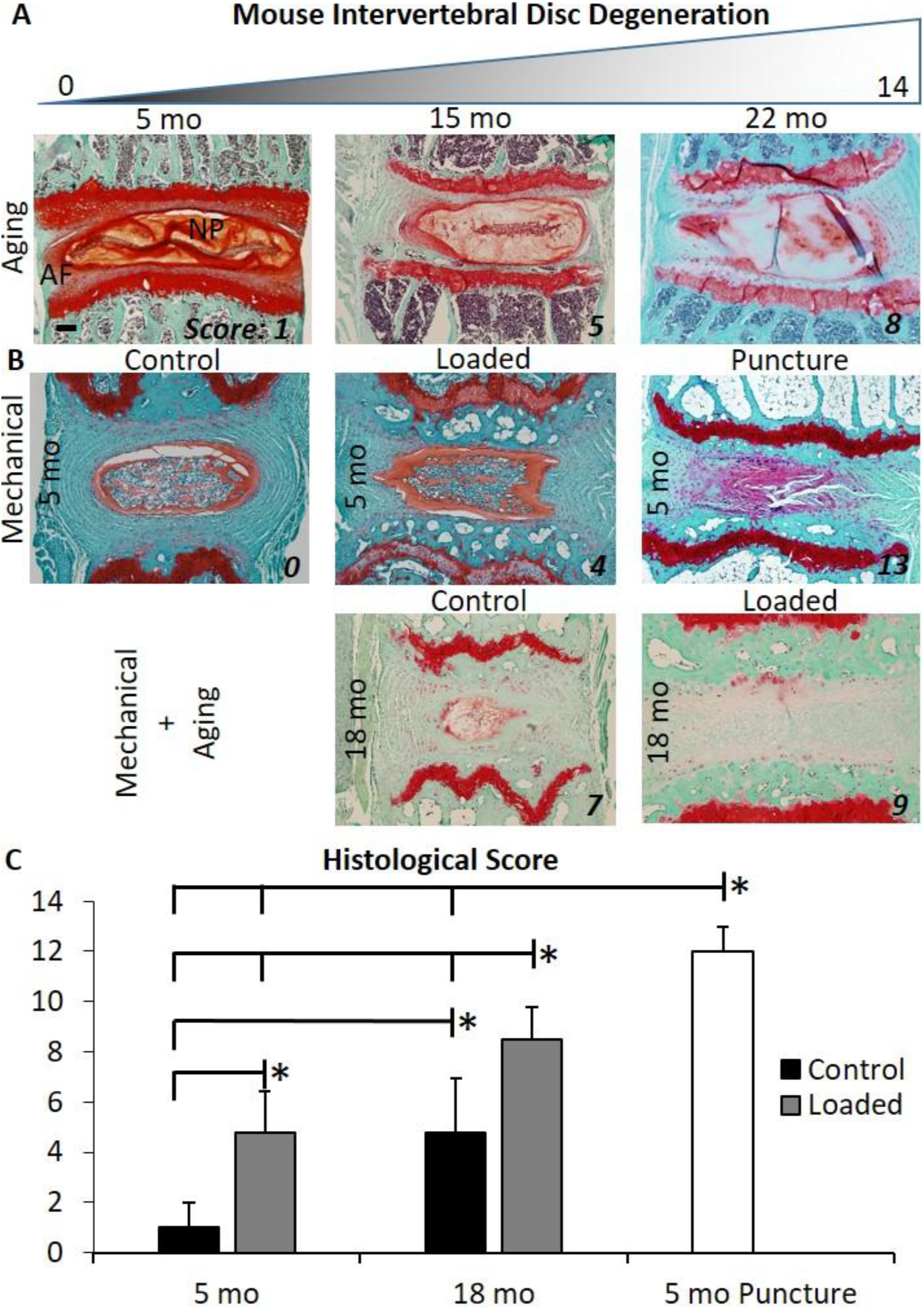
Mouse intervertebral disc (IVD) degeneration was scored histologically on a 0-14 scale. Individual scores are noted for each representative image. (A) Mouse intervertebral disc (IVD) degeneration increased with aging in the lumbar region. (B) Mechanical injury by tail compression or puncture induce IVD degeneration in 5 mo and 18 mo mice. (C) Quantification of the tail IVD degeneration. Scale bar: 100 μm. *: p<0.05.

### Osterix (OSX) expression and WNT signaling are suppressed by mechanical loading and aging

qPCR confirmed that aging and loading enhanced expression of catabolic and inflammatory markers and suppressed the expression of transcription factors and WNT signaling. Mechanical loading increased catabolic markers MMP3 and MMP13 by ≥7 fold in 5 mo IVD (**Fig. 2**). MMP3 and MMP13 were also upregulated by compression in aged mice, but MMP13 upregulation in middle-aged 12 mo IVD was less than in 5 mo IVD. MMP13 and ALPL are also markers of hypertrophic chondrocytes (D’Angelo et al., 2000) and aging reduced their expression by ≥50%. Secondly, IVD compression reduced the expression of key transcription factors OSX (Sp7) and Brachyury (T) by 50% and increased the gene expression of LAMIN-A, a marker of maturing differentiation (Constantinescu et al., 2006), by 1.5-fold. Aging reduced the gene expression of OSX and RUNX2, where RUNX2 was undetectable. However, the suppression of OSX by loading in 12 mo IVD trended to be less than the suppression in 5 mo IVD (interaction p=0.06). Lastly, loading trended to upregulate markers of WNT signaling in 5 mo IVD, but no response was noted in middle-age IVDs, other than the upregulation of β-Catenin in 12 mo IVD. Aging reduced the expression of LRP5 and LRP6 (**Fig. 1S**), and loading increased LRP5 and AXIN2 expression in 5 mo IVD. In addition, aging and compressive loading increased IL1b and TNF-a gene expression between 3- and 64-fold (**Fig. 1S**). These data suggest that, compared to young-adult IVD, the degenerative-response of older IVD to compression is associated with less chondrocyte-like expression and a disruption of the genes mediating chondrocyte accrual.

**Figure 2.**
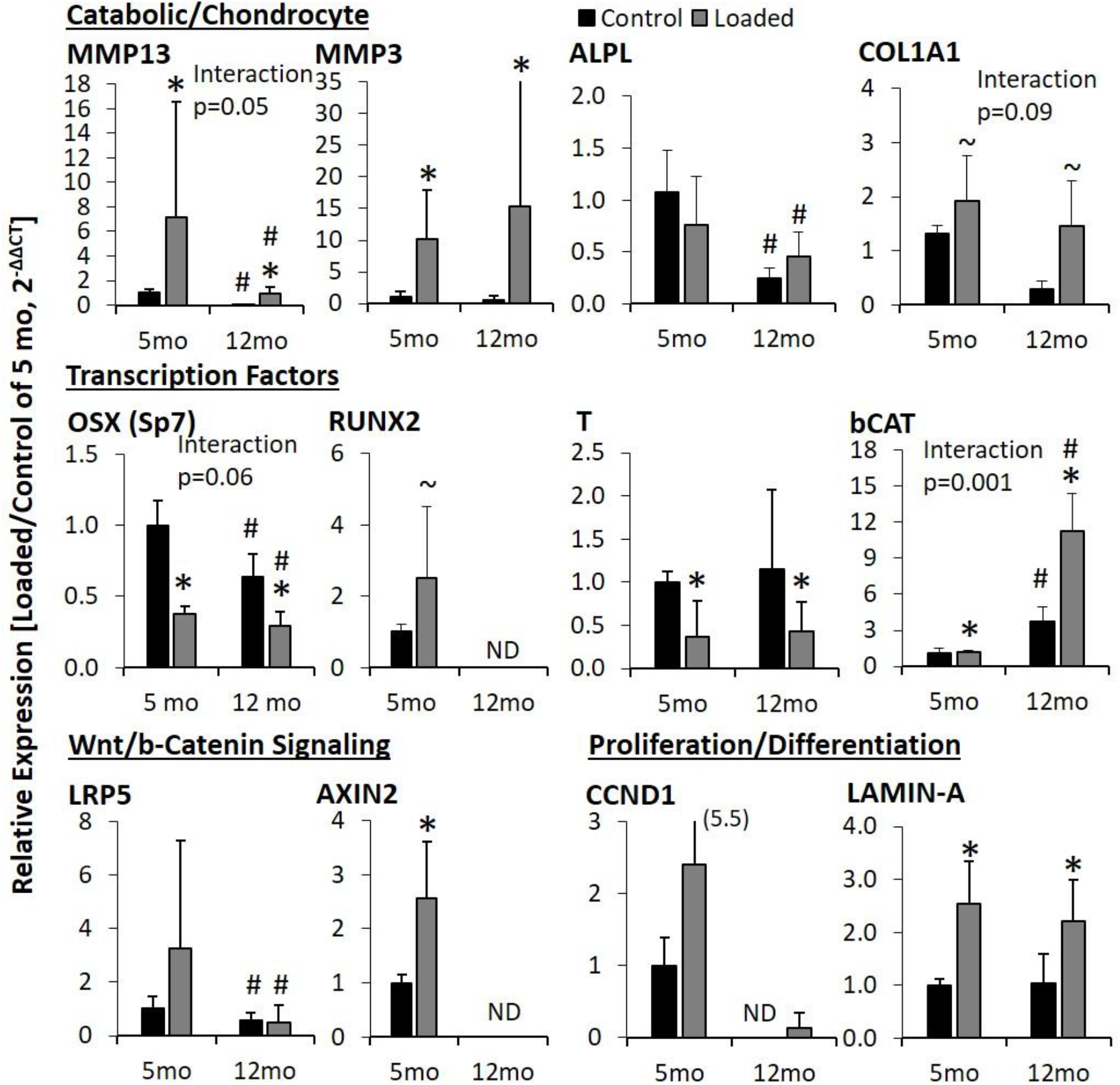
QPCR of 5 mo and 12 mo IVD subjected to tail compression. The relative gene expression in each loaded and control intervertebral disc was determined by normalizing to housekeeping gene IPO8 (CT value of 30) and then normalized to the control intervertebral disc. *: Main effect of loading, #: main effect of aging, p<0.05. ~:0<0.1

### Protein expression of osterix delineated a cell phenotype shift by mechanical overloading in the nucleus pulposus

In order to determine the location of the cells in the IVD expressing osterix and to corroborate the reduction of osterix with IVD degeneration, we stained for osterix protein in tail compressed IVD. First, osterix was expressed in osteoblasts of the cortical bone, trabecular bone, and endplate of the tail and lumbar vertebrae (**Fig. 3A**). In the tail and lumbar IVD, nucleus pulposus and outer annulus fibrosus cells expressed osterix. Because the suppression of osterix by mechanical compression was greatest in 5 mo IVD (**Fig. 2**), we determined the percentage of cells in the nucleus pulposus expressing high (small and dark), low (large and light) and no amount (no stain) of osterix protein in control and loaded IVDs (**Fig. 3C**). Mechanical loading reduced the percentage of cells in the nucleus pulposus with a small, darkly stained cell nucleus by 50% (**Fig. 3D**). The percentage of cells in the nucleus pulposus with a large, lightly stained nucleus did not change significantly, but was twice as many in loaded group. The net percentage of cells without osterix expression was unchanged and about 75% of the nucleus pulposus cells were positive for osterix. Osterix protein-intensity expression coincided with cytomorphology differences between notochordal and chondrocyte-like cells (Hunter et al., 2004; Smolders et al., 2012).

**Figure 3.**
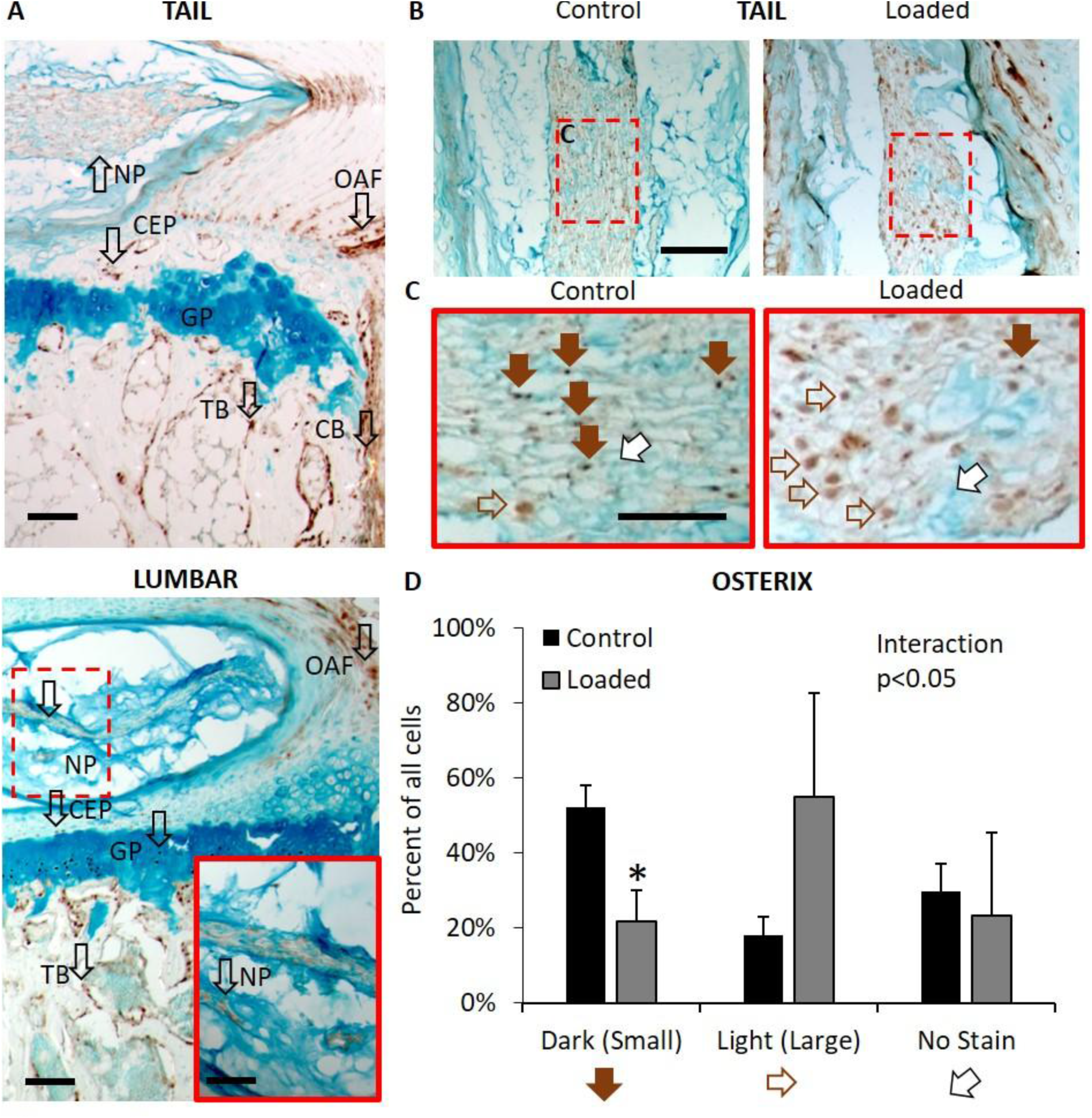
(A) Immunohistochemistry staining for osterix and counterstained with Safranin-O of tail and lumbar IVD. (A’) Magnification of the lumbar IVD. (B) Osterix staining of the NP of IVD subjected to tail compression. (C) Magnification of the NP and rotated by 90° clockwise. Solid brown arrows denote cells with small cell nuclei (relative to cell size) stained with a high-intensity of osterix expression (Dark), empty brown arrows denote cells with large cell nuclei stained with a low-intensity of osterix expression (Light) and white arrows denote cells with cell nuclei stained with no osterix expression (No Stain). Scale bar for A, B: 100 μm; A’:25 μm, and C: 12.5 μm. CB: cortical bone, CEP: cartilage endplate, GP: growth plate, NP: nucleus pulposus, OAF: outer annulus fibrosus, TB: trabecular bone. *: p<0.05.

### Osterix-expressing cells of the nucleus pulposus and outer annulus fibrosus decline with aging

Young-adult OsxCreERT2/Ai9-tdT tomato reporter mice corroborate the expression of osterix in the cells of the nucleus pulposus (45%) and the outer annulus fibrosus of tail IVD (**Fig. 4A, B**). Similarly to tail compression, aging reduced the expression of osterix in the cells of nucleus pulposus and outer annulus fibrosus. As expected, osteoblasts in the cartilage endplate, trabecular bone and cortical bones expressed osterix and the expression was not lost with aging. Little to no osterix expression was noted in the growth plate. Similarly, Ai9/tdT (without Cre) show little to no expression of the reporter (**Fig. 4C**).

**Figure 4.**
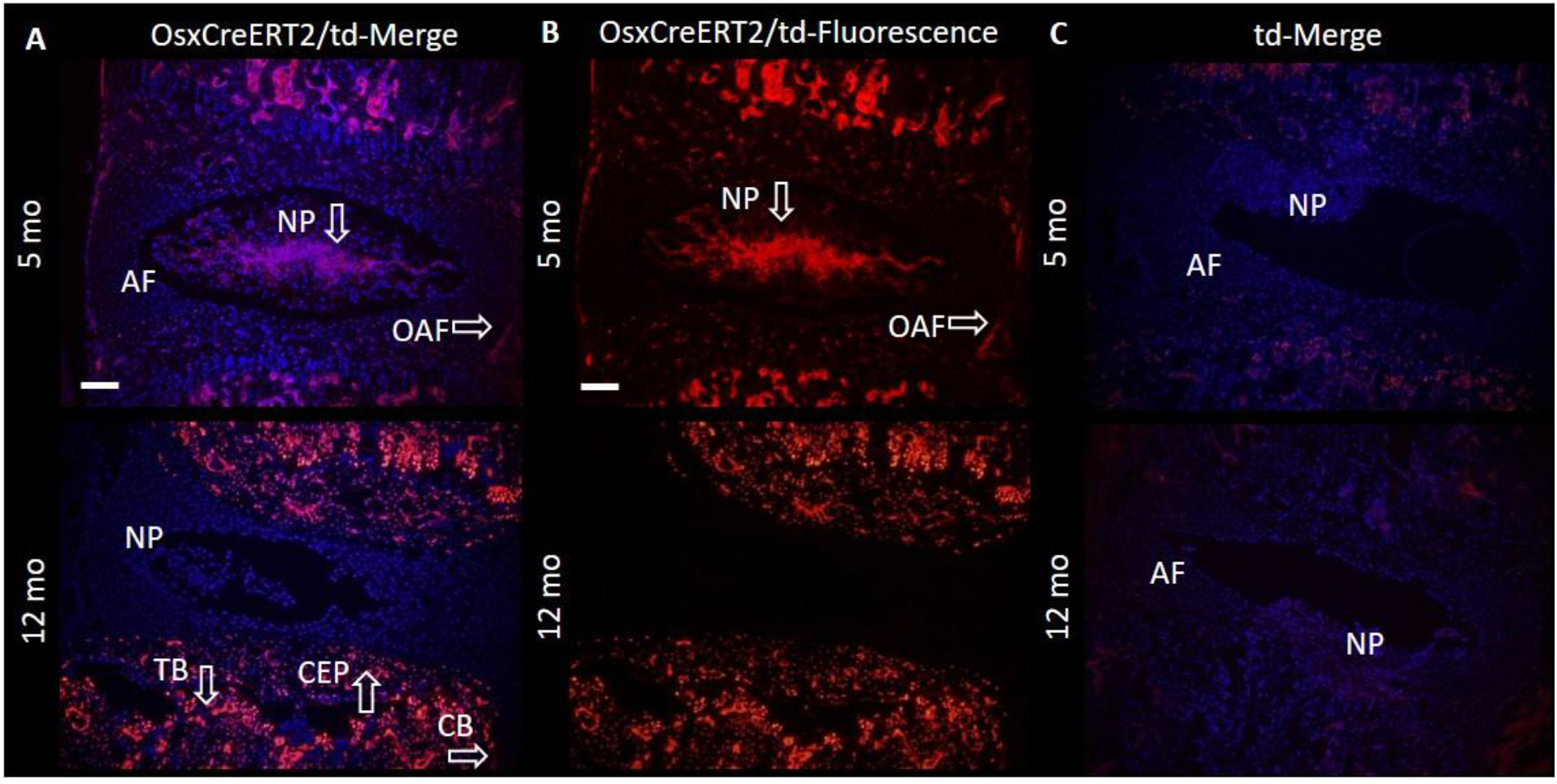
(A) Merged DAPI (blue) and immunofluorescence of osterix expression of tail IVD from 5 mo and 12 mo OsxCreERT2/tdT mice. Immunofluorescence of osterix expression of (B) IVD from 5 mo and 12 mo OsxCreERT2/td mice. (C) IVD of 5 and 12 mo td mice without OsxCreERT2. AF: annulus fibrosus, CB: cortical bone, CEP: cartilage endplate, NP: nucleus pulposus, OAF: outer annulus fibrosus, TB: trabecular bone. Scale bar: 100 μm.

### Osterix-specific deletion of LRP5 reduces WNT/β-Catenin signaling and mechanical properties and induces IVD degeneration

In order to mimic the inactivation of WNT signaling by aging and IVD degeneration, we targeted osterix-expressing cells to suppress LRP5. Therefore, we bred LRP5 cKO (LRP5 cKO: OsxCreER^T2^ mice/LRP5^fl/fl^/TOPGAL) and WT mice (LRP5^fl/fl^/TOPGAL). LRP5 deletion inactivated WNT signaling in lumbar and tail IVD (**Fig. 5A, E**). In lumbar IVD, the KO reduced WNT signaling in the nucleus pulposus by 95% and in the annulus fibrosus by 40%; retaining expression in the inner annulus fibrosus (**Fig. 5B**). In the tail IVD, the KO reduced WNT signaling in the nucleus pulposus by 60%, but did not change WNT signaling in the annulus fibrosus as none was detectable (**Fig. 5F**). Deficiency of LRP5 induced lumbar IVD degeneration originating from changes in the nucleus pulposus (**Fig. 5C, D**), whereas histological changes were not noted in tail IVD (**Fig. 5G, H**). Cell clusters, unclear demarcations between the nucleus pulposus and the annulus fibrosus, and large inner annulus fibrosus cells were the common degenerative features in the LRP5 cKO lumbar IVD.

**Figure 5.**
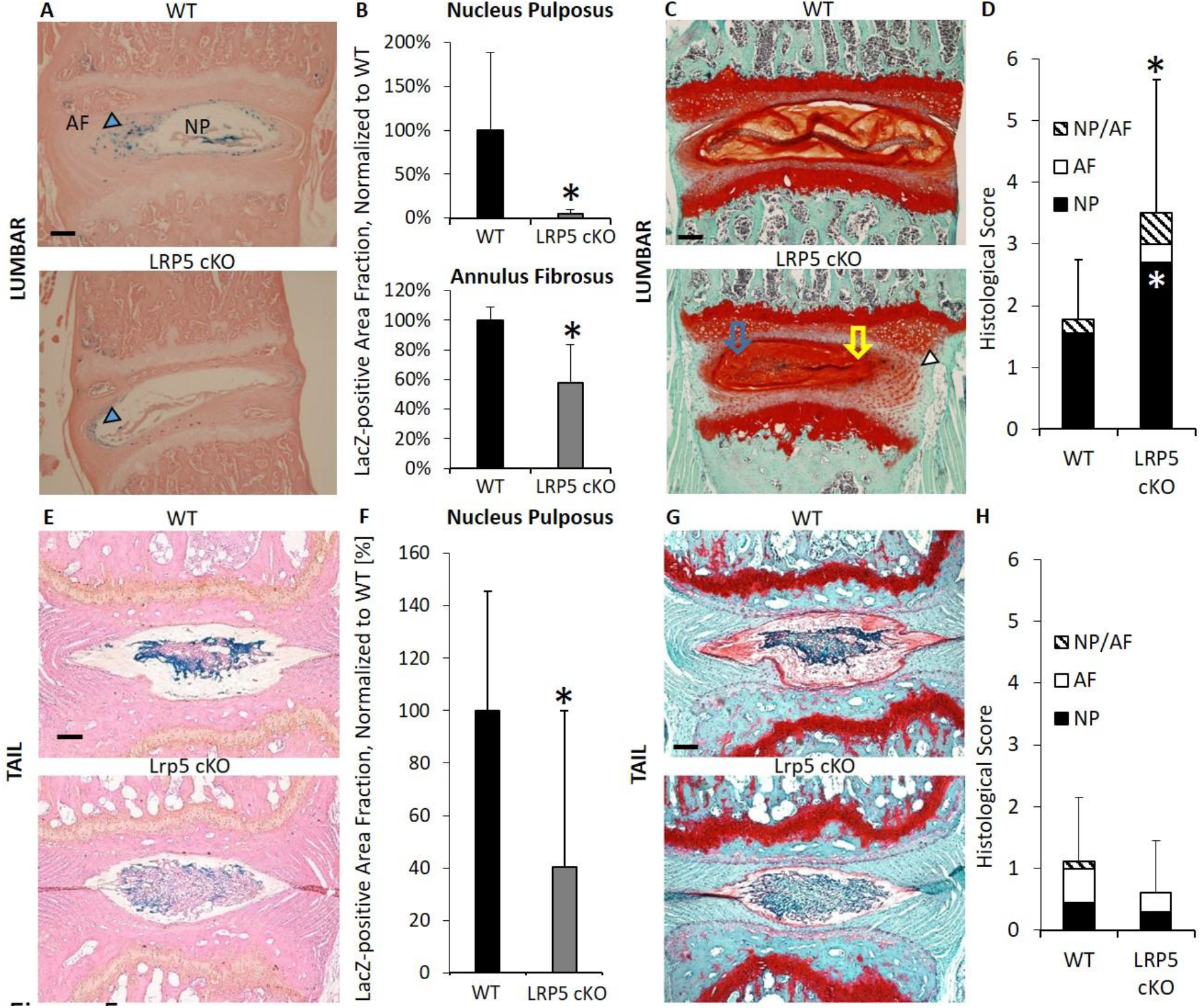
From WT and LRP5 cKO mice, (A) LacZ staining for WNT signaling (blue arrow head), quantification of WNT signaling, Safranin-O/Fast green staining and histological scoring of (A-D) lumbar and (E-H) tail IVD. Scale bar: 100 μm. *: p<0.05.

Deficiency of LRP5 in osterix-expressing cells reduced the mechanical properties of the IVD. Over the same range of motion (**Fig. 6A**), the structural stiffness of LRP5 cKO IVD was less than the control IVD by ≥45% (**Fig. 6B**). The morphology was not different between LRP5 cKO and WT IVD and, therefore, the material stiffness (modulus) was reduced in cKO similar to the structural property results (**Table 1**).

**Figure 6.**
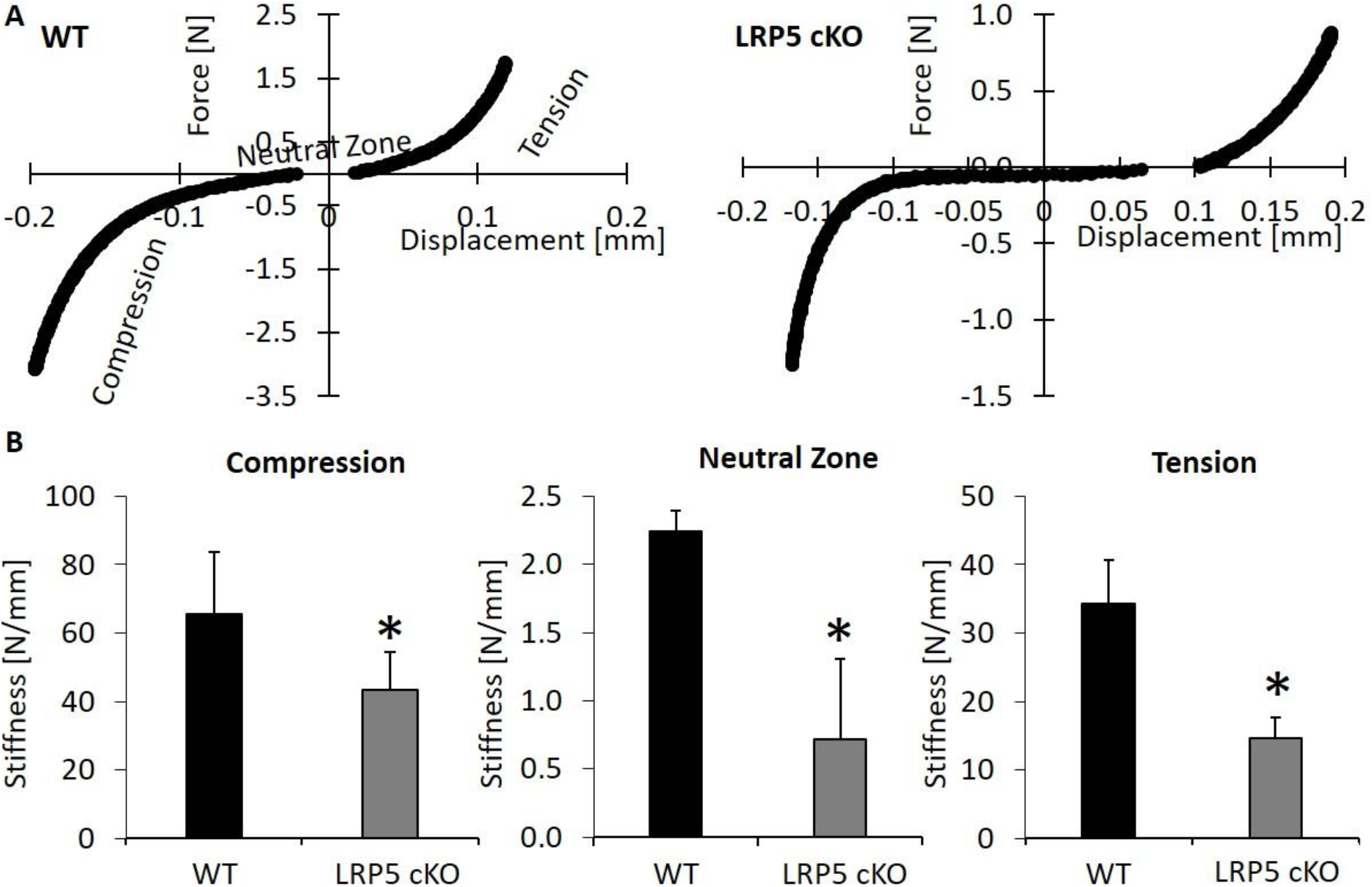
(A) Force/Displacement curves of control (WT) and LRP5 cKO tail IVD. (B) Stiffness in compression, neutral zone and tension of WT and LRP5 cKO IVD. *: p<0.05.

**Table 1.**
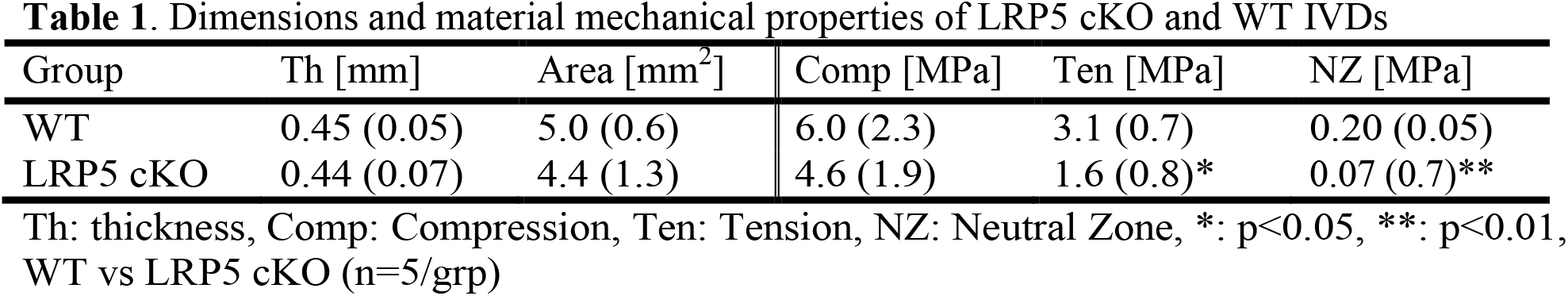
Dimensions and material mechanical properties of LRP5 cKO and WT IVDs

### LRP5-deficiency induces differential gene expression between lumbar and tail IVD

The gene expression of the lumbar IVD indicated a pattern consistent with IVD degeneration, where osterix, notochordal markers (KRT8 and KRT19) and chondrocyte marker MMP13 (D'Angelo et al., 2000) were downregulated by ≥50%, and aggrecan protease Adamts5 was upregulated by 60% (Fig. 7A). Contrarily, despite reduced mRNA osterix and β-Catenin expression (**Fig. 7B**), tail IVD had 2-fold more expression of notochordal markers, suggesting regeneration. We already confirmed that LRP5 cKO IVD had less WNT signaling than WT IVD, therefore we wanted to determine whether the remaining WNT signaling in the nucleus pulposus of KOs was due to inefficient targeting of osterix. Osterix (brown) and WNT signaling (blue) prominently co-stained in the WT (**Fig. 7C**). In contrast, in LRP5 cKO IVD, a majority of the osterix-positive cells appeared without WNT signaling and the cells that retained WNT signaling did not stain for osterix. Lastly, gene expression of WNT signaling-related genes was highly correlated to expression of the extracellular matrix and transcription factors. For instance (**Fig. 2S**), β-Catenin was associated with tensile stiffness (R^2^=0.60, p<0.05) and the mRNA expression of LRP5 (R^2^=0.78, p<0.05), and LRP6 was associated with aggrecan (R^2^=0.74, p<0.05) and RUNX2 (R^2^=0.85, p<0.01).

**Figure 7.**
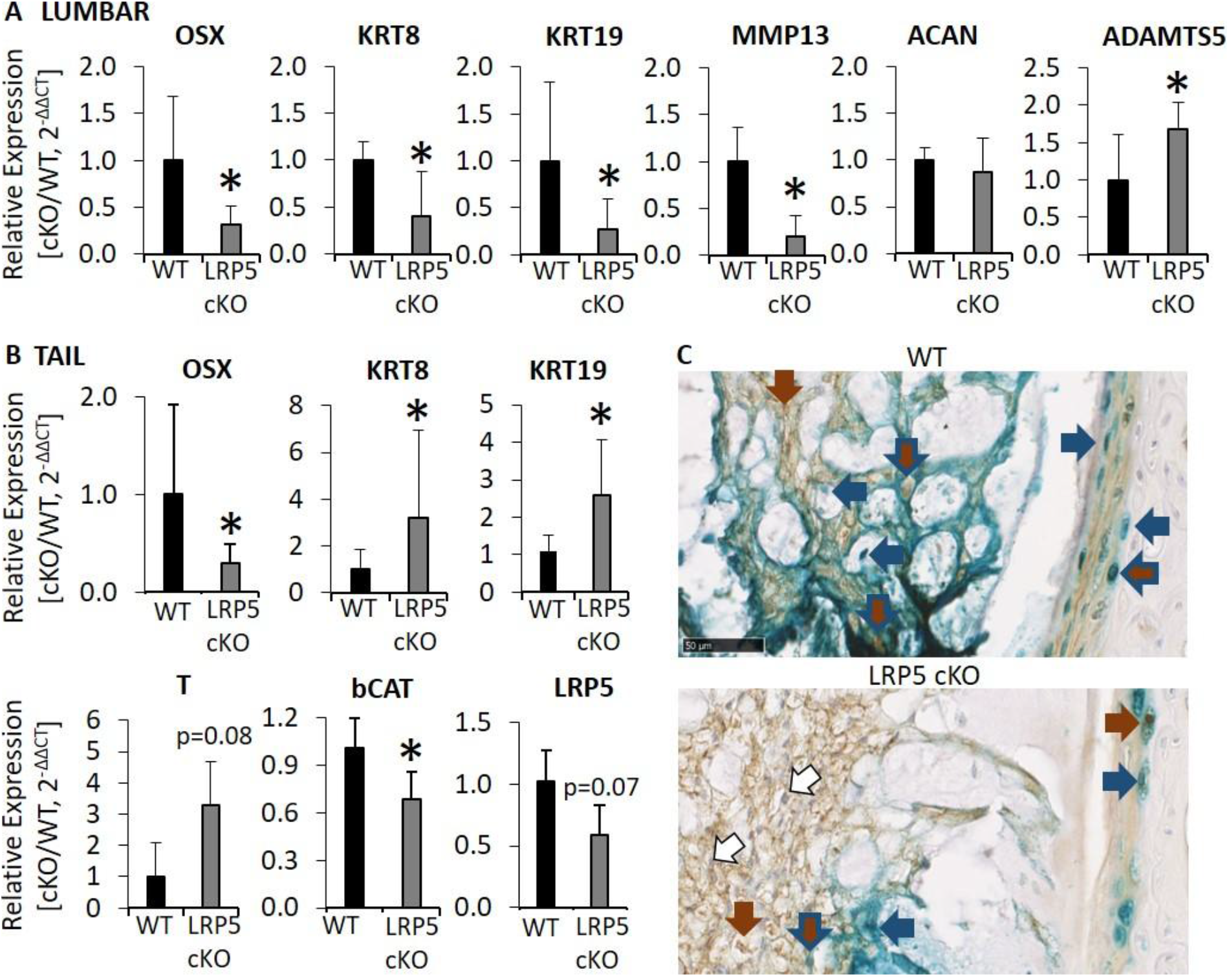
QPCR of (A) lumbar and (B) tail IVD from WT and LRP5 cKO mice. (C) LacZ staining for WNT signaling (blue arrow) and immunohistochemical staining for osterix (brown arrow) from WT and LRP5 cKO tail IVD. Cells stained for both are brown with blue lining. Sclae bar: 50 μm *: p<0.05.

## DISCUSSION

### Overview

We aimed to clarify the induction of chondrocyte-like cells with IVD degeneration by determining the changes in a key transcription factor of chondrogenesis (SP7/Osterix) during IVD degeneration and aging, and by mimicking age-related inactivation of WNT using an inducible, conditional KO. Both aging and mechanical compression reduce the expression of osterix, and loss of osterix was associated with cellular expression changes. IVD degeneration was heightened in old IVD following mechanical compression and regulation of transcription factors controlling chondrogenesis (Osterix and RUNX2) were impaired in old IVD. Lastly, WNT signaling in osterix-expressing cells of the IVD was reduced by deleting a WNT ligand receptor. These conditional KO IVD induced a level of IVD degeneration on par with aging and mechanical compression. Overall, these data implicate osterix as an important contributor to IVD regeneration.

### Osterix in Healthy IVD

The OsxCreERT2 mouse is commonly used to target bone cells in the osteoblastic lineage (Nakashima et al., 2002) and, while osterix is not expressed in growing mice (Zheng et al., 2019), we note that this inducible-Cre targets adult IVD cells in the nucleus pulposus and outer annulus fibrosus (Table 2). The OsxCreERT2 did not targeting cells in the inner annulus fibrosus as demonstrated by lack of expression of osterix by immunoflourescence and immunohistochemistry and by expression of WNT signaling in the inner annulus fibrosus following conditional deletion of LRP5 in osterix expressing cells. The utility of such a Cre has yet to be fully explored, but the only other inducible Cre drivers that target multiple regions of the IVD are the AcanCreERT2 (NP and AF) (Henry et al., 2009) and the Scleraxis-Cre (IAF and OAF) (Torre et al., 2019). The majority of the rest target one of the three regions of the IVD (Bedore et al., 2016; Chen et al., 2014b; Choi and Harfe, 2011; Imuta et al., 2013; McCann et al., 2012). The consequences of mutations in osterix on the IVD are unknown, but are associated with osteogenesis imperfect in global mutations (Lapunzina et al., 2010) and impaired chondrocyte differentiation when chondrocytes are targeted (Nishimura et al., 2012).

**Table 2.** Intervertebral disc-specific transgenic models with inducible Cre recombinase activity

| CreERT2 | Noto<br>(McCann<br>et al.,<br>2012) | Shh<br>(Choi<br>and<br>Harfe,<br>2011) | Foxa2<br>(Imuta<br>et al.,<br>2013) | Acan<br>(Henry<br>et al.,<br>2009) | Col2<br>(Chen<br>et al.,<br>2014b) | Scx<br>(Torre<br>et al.,<br>2019) | Col1<br>(Bedore<br>et al.,<br>2016) | Osx |
| --- | --- | --- | --- | --- | --- | --- | --- | --- |
| NP | + | + | + | + | - | - | - | + |
| IAF | - | - | - | + | + | + | - | - |
| OAF | - | - | - | + | - | + | + | + |

NP: Nucleus pulposus, IAF: Inner annulus fibrosus, OAF: Outer annulus fibrosus, Shh: Sonic hedgehog, Foxa2: Forkhead box a2, Acan: Aggreacan, Col2: Collagen type 2, Scx: Scleraxis, Col1: Collagen type 1, Osx: Osterix (Sp7), +: Present, -: Absent

### WNT Ligand Receptor LRP5 is Critical for WNT Signaling in IVD

Alongside loss of WNT signaling with aging and compression (Holguin et al., 2014; Holguin and Silva, 2018), cell membrane receptor for WNT ligands LRP5 (Gong et al., 2001) was reduced with aging and compression. Next, we showed that 1 month-deletion of LRP5 in osterix-expressing of young-adult mice impaired WNT signaling and induced IVD degeneration. These data suggest that WNT ligand(s) that bind to LRP5 alter canonical WNT signaling in the IVD. Our previous data show that compression reduces WNT signaling and ligands WNT16, 7b and 10a and, by contrast, stabilization of β-Catenin in the IVD elevates WNT16 among other genes (Holguin and Silva, 2018). WNT16 and WNT 7b alter canonical WNT signaling in other musculoskeletal tissues (Alam et al., 2016; Chen et al., 2014a; Nalesso et al., 2017) and may serve as targets for IVD therapy.

### IVD Degeneration Suppresses Osterix and Promotes Chondrocyte-like Cells

Here, we show that aging, mechanical compression, and inactivation of WNT signaling by an LRP5 cKO reduced the expression of osterix in the IVD and led to IVD degeneration. Immunohistochemical staining for osterix in the NP of compressed IVD noted a cell shift by the intensity of the staining and morphology of the cell. Tail compression is known to induce loss of notochordal cells in the NP prior to cell death (Yurube et al., 2014) and elevate chondrocyte-like cells. Currently, the role of osterix in the IVD is unclear but, in bone, osterix is a critical transcription factor that drives osteoblastic cell differentiation (RUNX2 before osterix and β-Catenin after osterix). Loss of osterix (Nakashima et al., 2002) or β-Catenin (Day et al., 2005) diverts differentiation from osteoblastogenesis to chondrogenesis. First, it is important to note that while chondrocytes are similar to the cells of IVD, the phenotypic expressions have some differences (Clouet et al., 2009). Nevertheless, our data agree with elevation of early chondrogenesis markers RUNX2 and β-Catenin (WNT signaling) during IVD degeneration (Iwata et al., 2015; Sato et al., 2008; Smolders et al., 2012; Wang et al., 2012). As such, in order to potentiate terminal differentiation towards hypertrophic chondrocytes, our data show osterix and WNT signaling declined in the compressed IVD of aged mice, which coincides with their known function in chondrogenesis (Ma et al., 2013) and previous data (Holguin and Silva, 2018). However, loss of osterix is in conflict with work that shows human IVD degeneration from Grade III to V is marked by calcification and elevated gene and protein expression of osterix (Shao et al., 2016). The models we used here represent early IVD degeneration and, therefore, osterix may be regulated differently in late-stage IVD degeneration.

### Aging Exacerbates Compression-Induced IVD Degeneration

Aging exacerbated the IVD degeneration induced by mechanical compression and was associated with loss of chondrocytic markers, transcription factors and WNT signaling. The upregulation of chondrocyte markers MMP13 (D’Angelo et al., 2000) and RUNX2 by mechanical compression in young-adult mice was subdued-to-absent in aged mice. Further, despite elevated mRNA and protein expression of β-Catenin by compression in aged mice and in patients with IVD degeneration (Holguin and Silva, 2018; Wang et al., 2012), WNT signaling is inactivated by compression in aged mice because of limited and impaired translocation of β-Catenin to the cell nucleus (Holguin and Silva, 2018; Simcha et al., 1998; Wu et al., 2019). Aging is a common factor of IVD degeneration that includes cell loss (Boos et al., 2002) and an accumulation of proliferation-inept cells that remain metabolically active. This aging-induced loss of NP cells may harm the IVD by reducing the notochordal cells that trigger the chondrocyte-like cells to stimulate glycosaminoglycan and aggrecan core protein synthesis (Aguiar et al., 1999) and by depleting the progenitors capable of cell replenishment (Sakai et al., 2012). Ultimately, cell loss may be the impetus for the recruitment of non-notochordal cells derived from the vasculature (Brisby et al., 2013) or possibly the inner annulus fibrosus (Merceron et al., 2014), which are less equipped to cope with the harsh mechanical, biochemical and oxygen-tension demands of the NP environment.

### Proposed Role of WNT signaling/Osterix in IVD Degeneration

We propose a model of 4 levels of NP health and the relationship between WNT signaling and osterix: (1) optimal regeneration, (2-Adult; 3-Aged) suboptimal regeneration and (4) degeneration (**Fig. 8A**). (1) During optimal regeneration, the notochordal cells with a high level of WNT signaling and no expression of osterix contribute to the regeneration of the IVD. We show here and in a previous study (Holguin and Silva, 2018), IVDs with elevated WNT signaling and notochordal expression were protected from IVD degeneration. During suboptimal regeneration, chondrocyte cells may contribute a greater role since notochordal cells without osterix are limited. (2) In this phase, WNT signaling stimulation of progenitors differentiate into notochordal cells with high-osterix/WNT signaling, which then differentiate into chondrocyte-like cells by suppression of WNT signaling and osterix (**Fig. 8B**). (3) Aging limits chondrocyte-like differentiation in two ways: (i) by suppression of progenitor differentiation into notochordal cells by limited WNT signaling and (ii) by suppression of notochordal cell differentiation to chondrocyte-like cells by limited downregulation of osterix. (4) Full IVD degeneration is characterized by loss of both notochordal and chondrocyte-like cells. Similar patterns of transcriptional regulation occur with osteoblastogenesis/chondrogenesis (Long, 2011; Ma et al., 2013).

**Figure 8.**
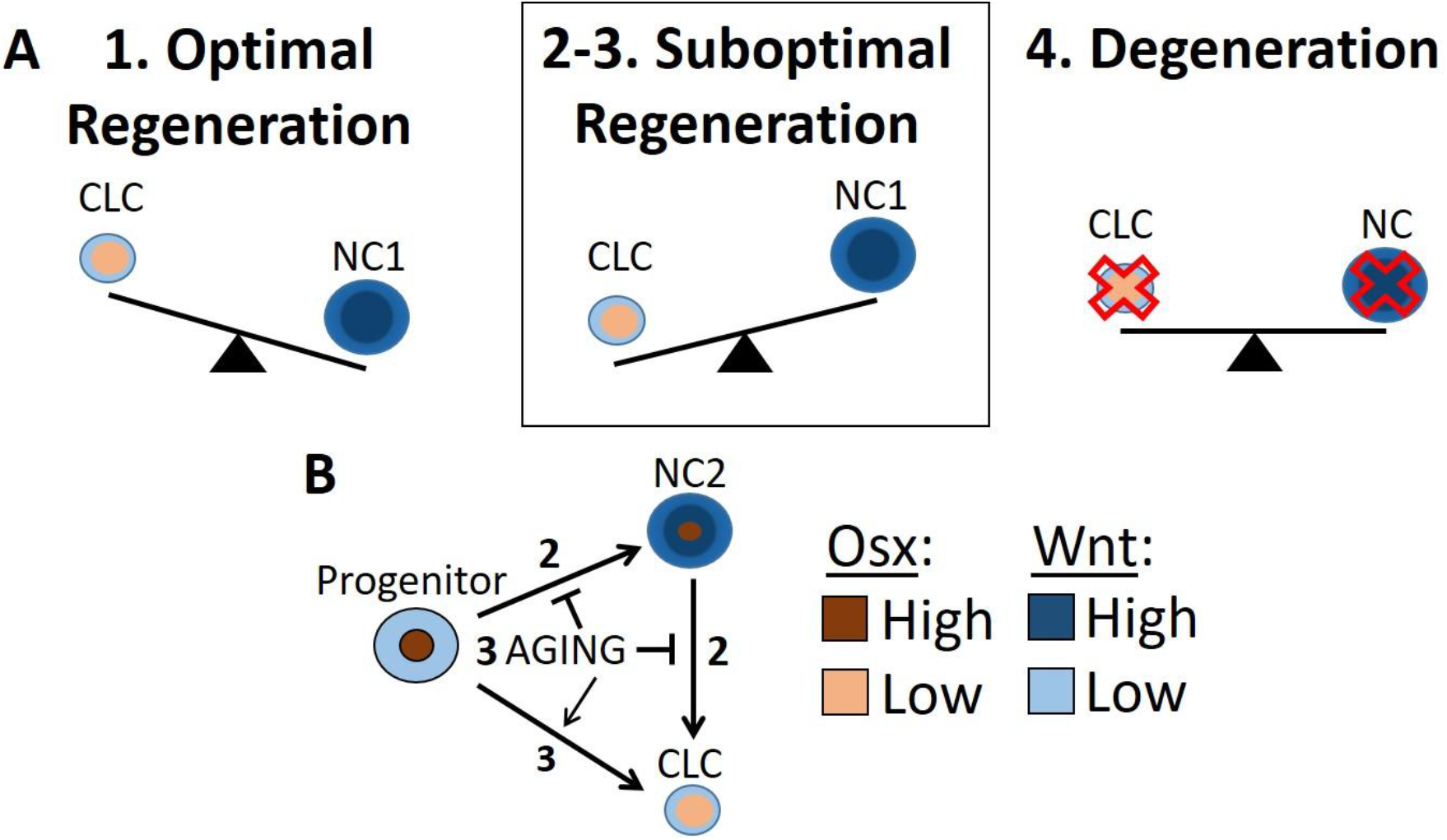
(A) Proposed role between chondrocyte-like (CLC) and notochordal (NC1) cells during optimal regeneration, suboptimal regeneration and IVD degeneration. NC1 cells have high WNT signaling and no osterix. IVD degeneration is characterized by a loss (X’s) of both cells. (B) During suboptimal regeneration, progenitor cells with high osterix expression gain WNT signaling to become NC2 and finally become CLC by losing osterix and WNT signaling. Aging limits differentiation of progenitors to CLC through NC2 and promotes it directly, which is suboptimal because fewer NC cells become involved.

### Limitations

All of the gene expression in the study was conducted in intact IVD, which does not separate changes between the nucleus pulposus and annulus fibrosus. For instance, gene expression of LRP5 was not significantly downregulated in the tail IVD of LRP5 cKO, but the functional reduction of Wnt signaling that is downstream of LRP5 using the TOPGAL transgene of the mouse corroborates that LRP5 was deleted. Secondly, it is unclear why IVD degeneration did not occur in the tail IVD of LRP5 cKO mice, but this lack of impact also occurs in the tail IVD of β-Catenin conditional KO mice when driven by a ShhCreERT2 (Holguin and Silva, 2018). We suspect that tail IVDs are less sensitive to change than lumbar IVDs because tail IVDs are not subjected to the same complexity and magnitude of mechanical stresses as lumbar IVD and therefore tail IVD degenerate more slowly with aging (Holguin et al., 2014).

## Conclusions

Osterix (SP7) is a critical transcription factor in osteogenic/chondrogenic differentiation but little is known of its role in the IVD. Young-adult, healthy IVD express osterix in the annulus fibrosus and nucleus pulposus, but lose osterix with advanced aging, IVD degeneration and targeted inactivation of WNT signaling. Overall, these data indicate that age-related inactivation of WNT signaling in osterix-expressing cells may limit regeneration by depleting progenitors and attenuating the expansion of chondrocyte-like cells.

## Acknowledgements

The authors gratefully acknowledge Keith Condon and Evan Buettmann for assistance with the histology for osterix and Jiannong Dai for assistance completing QPCR. This work was supported by NIH grants R01 AR047867 (MJS) and F32 AR064667 (NH), by the Washington University Musculoskeletal Research Center (P30 AR057235, T32 AR060719). Last, but not least, we would like to thank the Indiana Center for Musculoskeletal Health (ICMH) and Department of Mechanical and Energy Engineering for the start-up.

## Author Contributions

MJS: experimental design, data analysis, wrote manuscript; NH: Experimental design, data collection, data analysis, wrote manuscript.

## SUPPLEMENTAL FIGURES

**Figure 1S.**
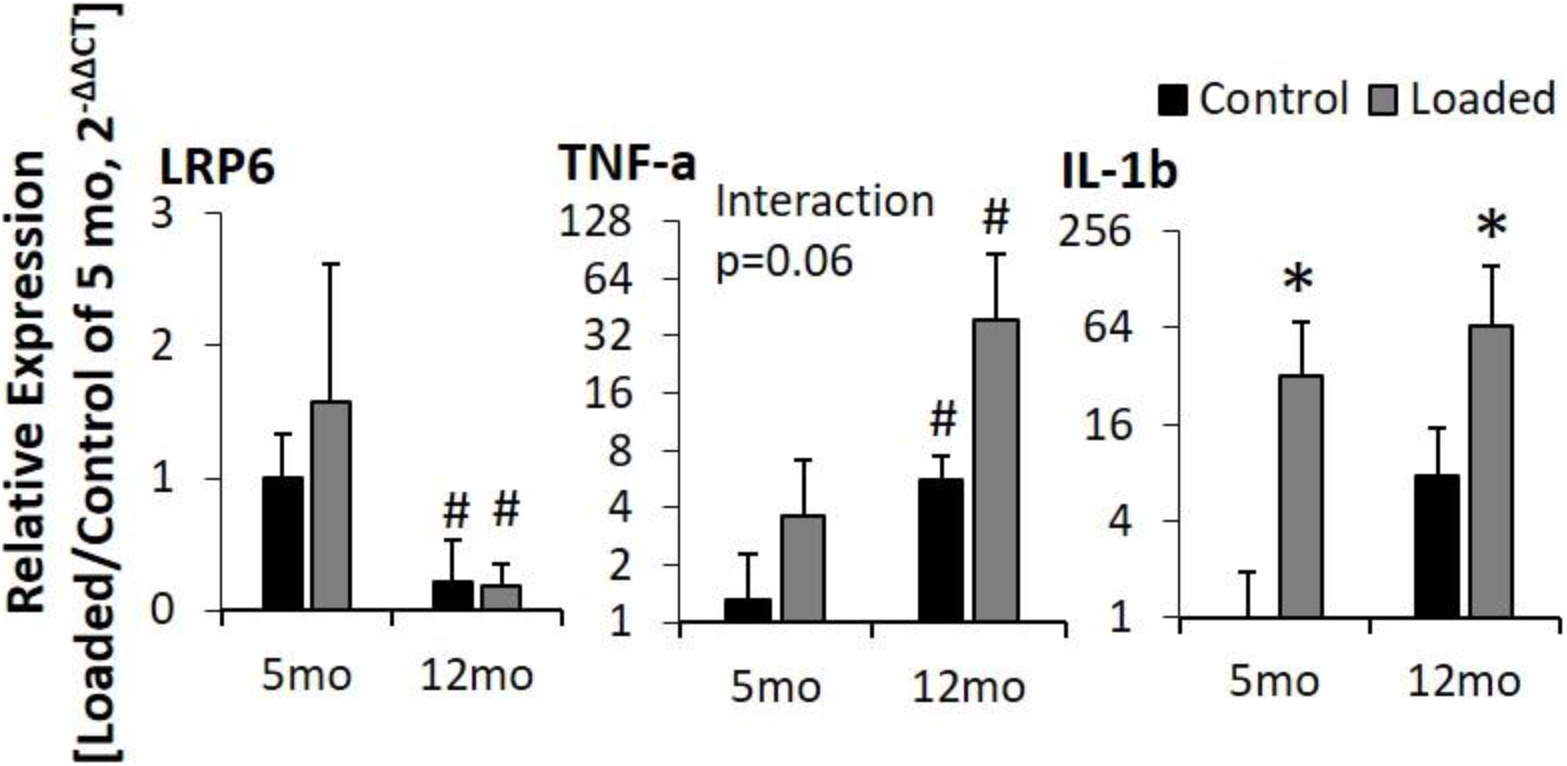

**Figure 2S.**
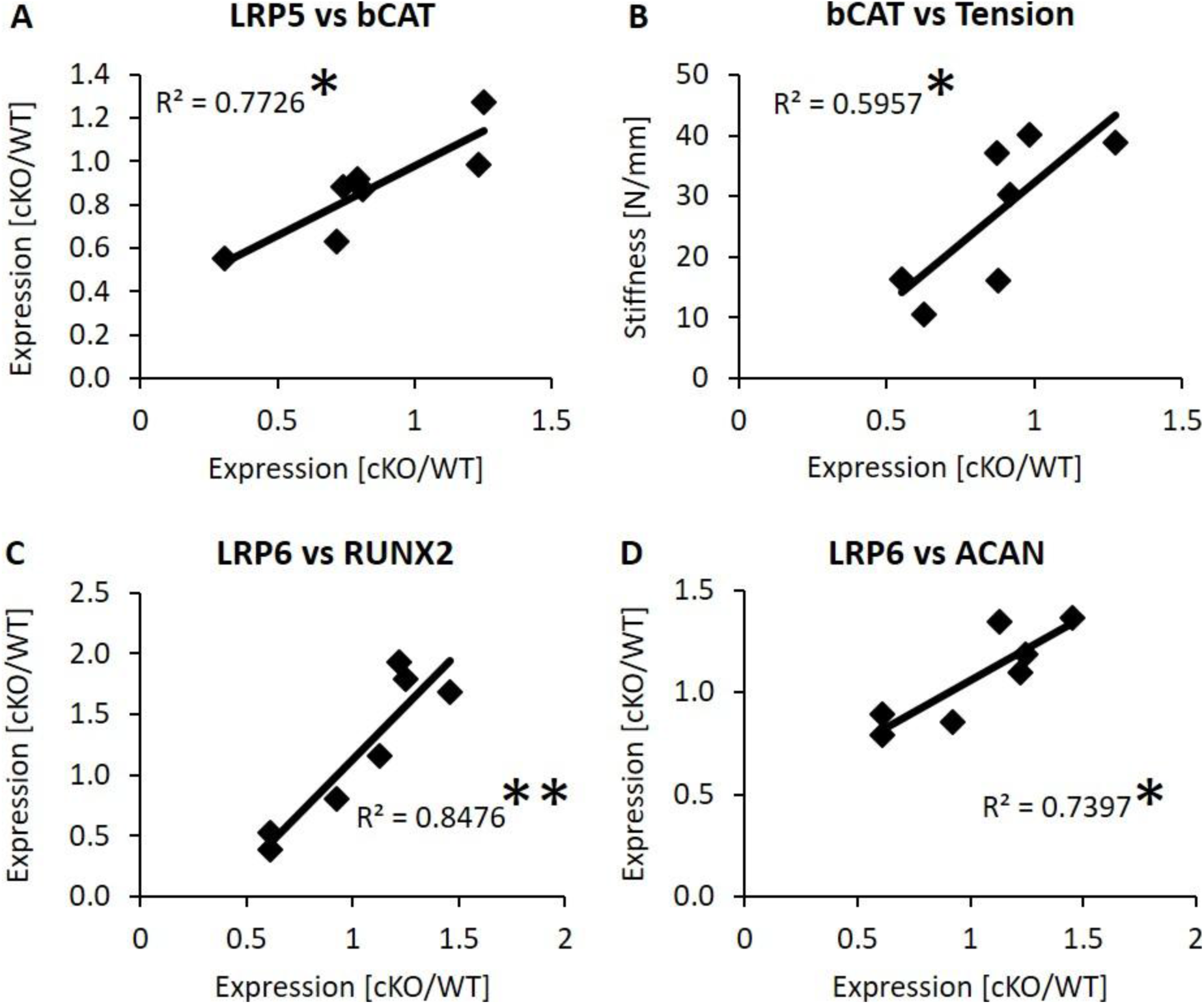

